# Objective (Real-world) Motion Responses in Scene Responsive Regions

**DOI:** 10.1101/145573

**Authors:** Didem Korkmaz Hacialihafiz, Andreas Bartels

**Affiliations:** Centre for Integrative Neuroscience, University of Tübingen, 72076, Tübingen, Germany; Max Planck Institute for Biological Cybernetics, 72076, Tübingen, Germany; Department of Psychology, University of Tübingen, 72076, Tübingen, Germany

## Abstract

We perceive scenes as stable even when eye movements induce retinal motion, for example during pursuit of a moving object. Mechanisms mediating perceptual stability have primarily been examined in motion regions of the dorsal visual pathway. Here we examined whether motion responses in human scene regions are encoded in eye- or world centered reference frames. We recorded brain responses in human participants using fMRI while they performed a well-controlled visual pursuit paradigm previously used to examine dorsal motion regions. In addition, we examined effects of content by using either natural scenes or their Fourier scrambles. We found that parahippocampal place area (PPA) responded to motion only in world- but not in eye-centered coordinates, regardless of scene content. The occipital place area (OPA) responded to both, objective and retinal motion equally, and retrosplenial cortex (RSC) had no motion responses but responded to pursuit. Only PPA’s objective motion responses were higher during scenes than scrambled images, although there was a similar trend in OPA. These results indicate a special role of PPA in representing its content in real-world coordinates. Our results question a strict subdivision of dorsal “what” and ventral “where” streams, and suggest a role of PPA in contributing to perceptual stability.

## Introduction

Keeping a stable visual perception is one of the crucial roles of the visual system. Our visual system continuously integrates retinal inputs with eye-, head- and body movements in order to keep our subjective visual percept stable and in register with the external world. The mechanisms behind this constant updating are only partially known. Prior studies have primarily focused on high-level parietal regions, identifying several that engage in so-called remapping, potentially allowing for stable perception (Fischer et al., 2012a, b; Galletti et al., 1990; Ilg et al., 2004; Zhang et al., 2004). Given that the content that is perceived as stable during eye movements typically consists of real-world scenes, it seems reasonable to examine this question also in scene-selective regions.

Scene processing has been shown to take place in PPA (Aguirre et al., 1998; Epstein and Kanwisher, 1998), RSC (Maguire, 2001) and OPA (also known as transverse occipital sulcus (TOS)) (Dilks et al., 2013; Grill-Spector, 2003; Hasson et al., 2003; Nakamura et al., 2000). Several studies have examined spatial updating in these regions using saccadic eye movements (Golomb et al., 2011; Ward et al., 2010) or snapshots of different viewpoints (Epstein et al., 2003; Epstein et al., 2005; Park and Chun, 2009; Sulpizio et al., 2014; Sulpizio et al., 2013). These studies have however not led to a clear agreement with regard to the reference frames encoded in scene regions. One study suggested that PPA and OPA utilize eye-centered coding while RSC does not show any preference (Ward et al., 2010). Another study found that PPA partially adapted to views of the same scene during saccadic eye movements, but this adaptation did not differ when the scene-snapshot moved with the saccade or not (Golomb et al., 2011). This study hence suggested that scene encoding is primarily eye-centered, yet with limited world-centered contribution. A problem with these saccade studies was that due to methodological considerations, saccades were executed on a blank screen, hence preventing true spatial updating of the scene during the saccade.

Here, we re-examine the important question whether scene regions encode visual input in eye-centered or world-centered coordinates using continuous motion and visual pursuit instead of using snapshots and saccades. This has multiple advantages. Pursuit can be carried out on the scenes rather than on intermittent blank screens, and updating occurs continuously rather than only a few times per stimulus block or between blocks. Updating-related signal can hence be expected to be considerably higher as it is generated continuously throughout each block. PPA and OPA have previously been shown to be motion responsive (Korkmaz Hacialihafiz and Bartels, 2015), as well as neurons of the parahippocampal gyrus in monkey (Sato and Nakamura, 2003). However, no study differentiated between retinal and world centered reference frames of these motion responses. Even among dorsal motion-selective regions only a subset encodes motion primarily in world-centered reference frame, such as V3A and V6 (Fischer et al., 2012a).

We designed stimuli according to a well-controlled 2x2 factorial design with the factors objective motion (on/off), pursuit (on/off) that allows separating eye- from world-centered motion encoding (Fischer et al., 2012a). A third factor of scene content (gray scale landscape and cityscape scenes or Fourier their scrambles (Korkmaz Hacialihafiz and Bartels, 2015)) was added to examine content-dependence of reference frame preference. To balance attention across all conditions, participants performed a central character-matching task at all times. Importantly, effects of eye movements cancelled out for the important contrasts, since conditions including pursuit eye movements were present in both sides of the equation. Scene responsive regions were identified using an independent localizer scan. We performed GLM whole-brain analyses as well as region of interest (ROI) analyses. Results show that the key scene responsive regions, PPA, OPA and RSC, can be completely dissociated on the basis of responsiveness to pursuit or their preference for the distinct reference frames.

## Materials and Methods

### Participants

17 healthy participants with normal or corrected-to normal vision (9 female, 1 left handed, age between 20 and 36, mean = 27.8 years) took part in this study after giving written informed consent. The study was approved by the ethics committee of the University Hospital of Tübingen.

### Experimental Setup

This study consisted of one main experiment, one functional localizer and one structural scan. The functional localizer aimed to identify scene- responsive regions: PPA, RSC, and OPA.

Visual stimuli were gamma corrected and back-projected onto a screen via a projector outside the scanner room. The screen was viewed via an angled mirror and subtended a visual field of 19 x 15 visual degrees.

The experiment was programmed using Psychtoolbox-3 (Brainard 1997, Kleiner, Brainard et al. 2007) on MATLAB 7.10.0 (The Mathworks, Natick, MA, 2010) and was presented using a windows PC.

### Main Experiment

Figure 1 illustrates the eight conditions of the main experiment. It was a 2 x 2 x 2 factorial design with the factors objective motion (on/off), pursuit (on/off) and scene (on/off). The design of the first two factors (on-screen motion and pursuit) was identical to that described in a prior study (Fischer et al., 2012a). Here, a background image (described by the third factor: either a natural scene or Fourier-scramble thereof) was either stationary or moved on a horizontal trajectory left- and right-wards. A horizontal trajectory was chosen as this corresponds to common eye- and head-rotations in natural situations and correspondingly predominates in feature movies. The velocity followed a sine function with a cycle of 3 s, and each block contained 4 cycles, hence lasting 12 s. The velocity varied between 0 and 3.08 deg/s, yielding a mean velocity of 2.53 deg/s. The motion extended 1.98 visual degrees in each direction. The starting direction of motion of each block was pseudorandomized and counterbalanced across runs.

**Figure 1.**
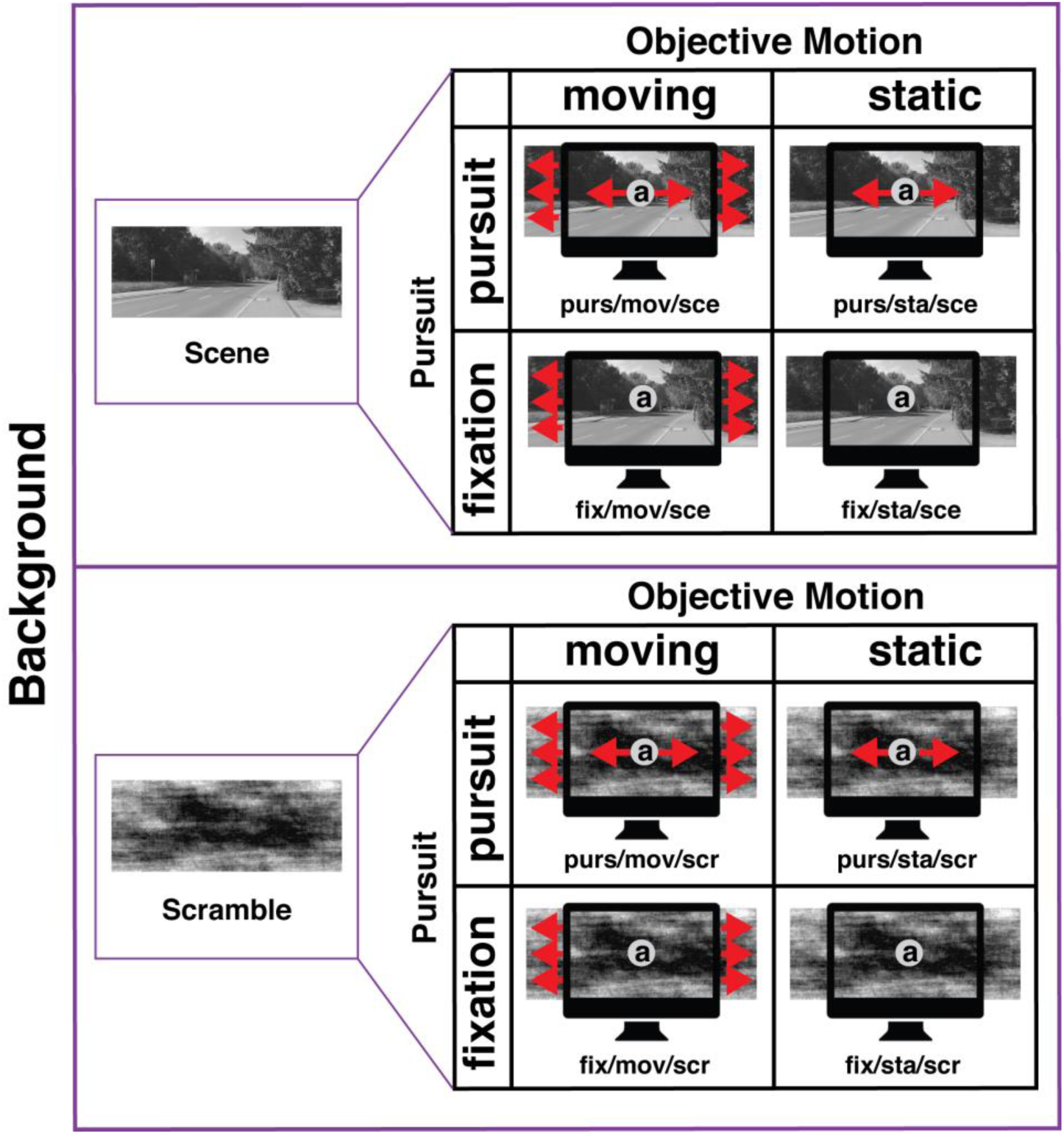
Conditions presented in the main experiment. A 2 x 2 x 2 factorial design with factors objective motion (moving, still), pursuit (pursuit, fixation) and scene (scene, scramble) resulted in eight conditions shown above. Objective motion was horizontal motion of the background image (scenes or scrambled images) and pursuit was horizontal motion of the fixation disk. There was a 1-back character-matching task inside the fixation disk (shown larger for illustration).

The same parameters applied to the fixation disc, that could also either be stationary or moving. In conditions when both, the fixation disc and the background moved, the motion of both was locked, such that there was no relative motion.

#### Background images

We picked 32 images of outdoor scenes (both landscapes and cityscapes) and converted them to gray scale. These gray scale images and their phase-scrambled versions composed the stimuli. In order to prevent unbalanced stimuli due to horizontal inequalities in the images, half of the images were left right flipped duplicates of the other half. All images were adjusted so that they had equal contrast and luminance (luminance: 144 cd/m2, contrast: 32.4 cd/m2 root-mean-square (RMS) contrast, resulting in an average Michelson contrast of 0.9004 ± 0.0925). Images were larger than the screen to allow their displacement while filling the screen at all times.

Phase-scrambled versions of the images were created using Fourier transformation and reconstruction with random phases. This resulted in preservation of low-level features of the image such as luminance, contrast and spatial frequencies while removing scene content. The same images were used in one of our prior studies (Korkmaz Hacialihafiz and Bartels, 2015).

#### Paradigm

The stimuli were presented in a block design. Each run consisted of 33 stimulus blocks. Each block lasted 12 seconds. The eight conditions were pseudorandomized and back matched so that each condition was preceded by all conditions with equal frequency across two runs. Each participant took part in 4 runs in total. Moreover, one additional block was added to the beginning of each run in order to ensure full counterbalancing for the first block. Each condition was presented 4 times in total in each run. Background images were randomly chosen for each block and only one image was used for an entire block.

Each run started with 6.9 seconds of gray screen with fixation and ended with 10 seconds of gray screen with fixation (luminance of gray screens: 144 cd/m2) leading to a total duration of 412.9 seconds. During the experiment, there was a gray fixation disk (width: 0.74 deg, luminance: 282 cd/m2) present at all times on the center of the screen, with the fixation task described below.

#### Fixation Task

In order to ensure fixation and balanced attention, participants were required to perform a 1-back character-matching task. The task was as follows: on the fixation disk a randomly chosen alphabetical character (a-z) was presented for 1 second each with 83 ms blank intervals in between. At random intervals, every 3 to 8 presentations, a repetition of the presented character occurred, which participants were required to report via button press. The timings of button presses were recorded and included in the GLM analyses as a regressor of no interest.

### Functional Localizer and ROI definition

We used a separate localizer experiment in order to localize scene selective regions PPA (Aguirre et al., 1998; Epstein and Kanwisher, 1998), RSC (Maguire, 2001) and OPA/TOS (Dilks et al., 2013; Grill-Spector, 2003; Hasson et al., 2003; Nakamura et al., 2000) for every subject in each hemisphere. The localizer consisted of 3 conditions; gray scale images of scenes (which were different than the ones used in the main experiment), faces and phase-scrambled versions of these scenes and faces. The localizer consisted of one run, and the stimuli were shown in a block design. In each block 5 different images from the same category were shown for 3 s each, yielding a block length of 15 s and 1 s gray screen following each block. The experiment started with 6.9 s of gray screen with fixation, had 27 blocks in total (9 times x 3 conditions) and ended with 10 s of gray screen with fixation. There was a fixation cross and participants were asked to fixate at this central fixation cross at all times. All three ROIs were identified using the contrast (scenes > faces), using the MarsBaR toolbox (Brett et al., 2002). In order to keep the ROIs similar in size across participants, we used an individual p-value for each participant and ROI when defining the ROIs (Fox et al., 2009; Murray and Wojciulik, 2004). Out of a total of 34 hemispheres, PPA was defined in 34 hemispheres, OPA in 28, and RSC in 33 hemispheres.

### Data Acquisition

T2* weighted functional images were acquired using a 64-channel phased-array head coil in a Siemens Magnetom PRISMA 3T scanner (Siemens, Erlangen, Germany) with the following parameters: voxel size 3 x 3 x 3 mm^3^, TR: 2.3 seconds, TE: 35 milliseconds, flip angle was 79°, 32 slices acquired in ascending order. In order to allow T1 equilibration, the first 3 volumes of data were discarded. Anatomical images were collected for each participant using T1-weighted images (1 x 1 x 1 mm^3^ resolution).

### FMRI Data Preprocessing and Statistical Analysis

Preprocessing was performed using SPM5 (www.fil.ion.ucl.ac.uk/spm) with the following steps: slice-time correction, realignment for motion correction, coregistration of the structural image to the mean functional image, normalization of the data to the SPM template in Montreal neurological institute (MNI) space, and spatial smoothing with 6 mm full-width at half maximum Gaussian kernel for single participants and 12 mm for group level analyses, respectively.

Data of each participant were analysed separately using the GLM (general linear model) in SPM5. We modelled each of the eight conditions as boxcars convolved by the canonical hemodynamic response function (hrf). Button presses were modelled as events. A total of seven regressors of no interest were included, consisting of six motion realignment regressors and one additional regressor for global signal variance (Desjardins et al., 2001; Van Dijk et al., 2010). The global signal variance regressor was orthogonalized to the conditions of interest. The data were high pass filtered using a cut-off value of 128 s. Beta images from the first level GLMs of each participant were used for group level analyses.

The ROI analyses were done by first extracting mean beta values for each ROI for each condition of each participant. Beta values were normalized in the range between 0 and 1 for each ROI and participant separately as follows: for each ROI of a given participant there were 32 mean beta values resulting from 4 runs and 8 conditions. The minimum of all 32 beta values was subtracted, and then all 32 beta values were divided by their maximal value. After this, an average beta value was calculated for each condition. This normalization ensured that all ROIs were comparable in mean and range of beta values. Repeated measures ANOVAs, as well as paired t-tests were conducted in order to analyse the effects of conditions using statistical analysis software IBM SPSS Statistics version 22.0. Greenhouse-Geisser correction was utilized in case of violation of sphericity as determined by Mauchly’s sphericity test.

The contrasts used in the analysis were defined as follows: “objective motion” was defined using all conditions with moving background versus all conditions with still background, i.e. left vs. right column in Figure 1. “Retinal motion” was defined by all conditions where retinal input changed versus all conditions where retinal input remained the same, i.e. in Figure 1 (purs/sta) + (fix/mov) versus (purs/mov) + (fix/sta). Objective versus retinal motion was hence equivalent to (purs/mov) versus (purs/sta) (Fischer et al., 2012a). The contrast “Scene” was defined using all conditions with scene vs. all with scramble. “Pursuit” was defined as all conditions with pursuit versus all conditions with eyes fixed (fixation). Note that all contrasts except for “pursuit” were balanced in terms of eye movements. The contrast “pursuit” contained both, eye-movement-related effects as well as effects related to peripheral visual stimulation induced by pursuit, but can nevertheless serve to functionally distinguish ROI properties.

### Eye Tracking

Eye tracking of participants during the main experiment was done using an infrared camera based eye tracker system (Eye-Trac 6; Applied Science Laboratories). Preprocessing included blink removal, and smoothing of x and y positions using a running average window of 200 milliseconds. We calculated the fixation accuracy by the root mean square error of actual eye position relative to the fixation disk for each condition across participants and runs. We then used repeated measures ANOVAs and t-tests to examine condition-related effects.

## Results

We investigated responses of independently localized scene-selective regions PPA, RSC and OPA to visual motion in world-centered (objective motion) and eye-centered (retinal motion) reference frames. The experiment was a 2x2x2 design with the factors pursuit (on, off) and visual motion (on, off), carried out using two types of content (natural scenes, their Fourier scrambles) as a third factor (see Figure 1). Figures 2A and 2B show raw and normalized mean beta responses to all conditions and each ROI, respectively. The contrasts of interest, such as for objective motion, retinal motion, or pursuit, were calculated using normalized responses that allowed for inter-region comparisons.

**Figure 2.**
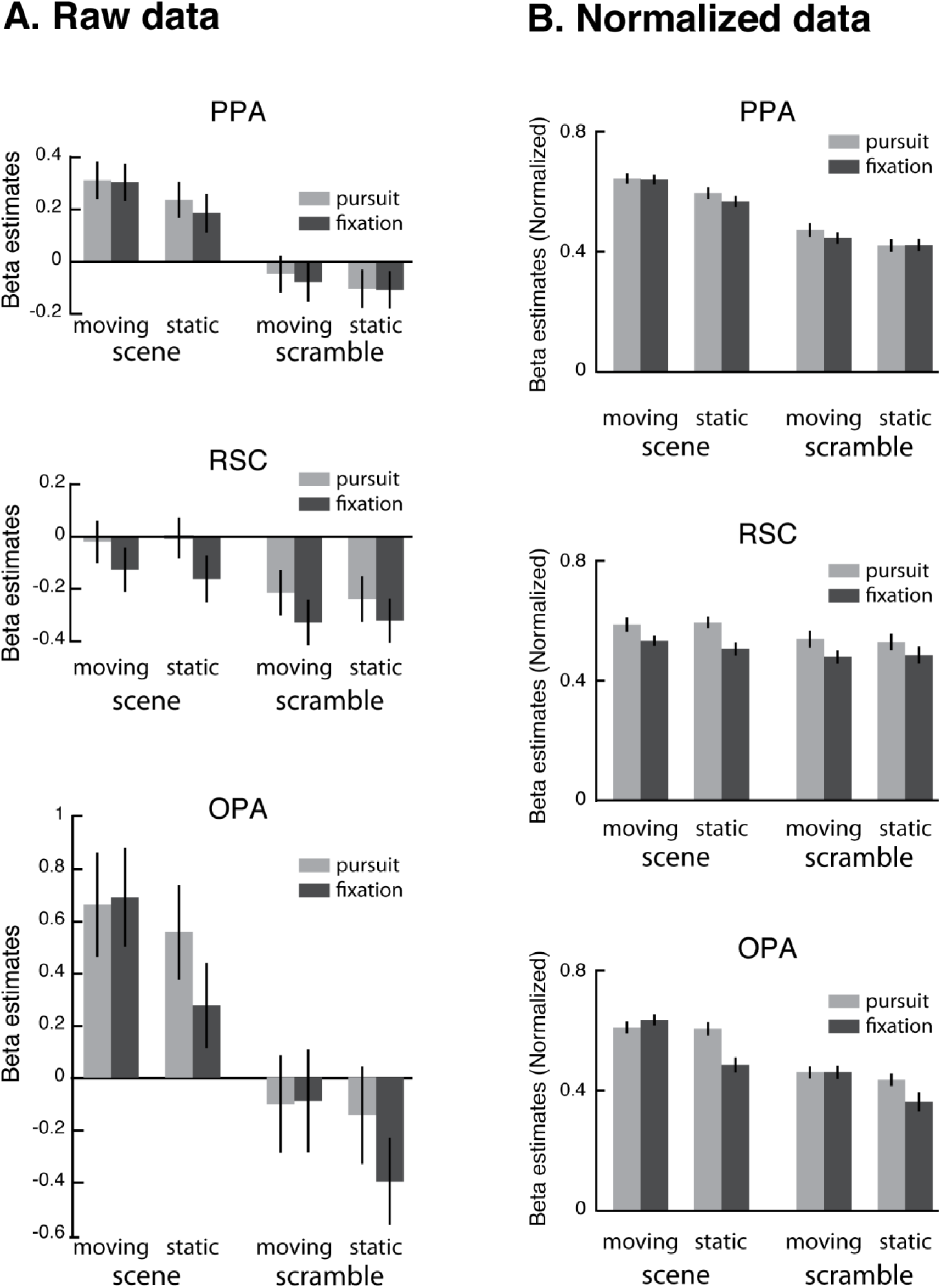
Responses to all eight conditions across ROIs. (A) Raw beta estimates in PPA, RSC and OPA. (B) Normalized beta estimates (see methods). Plots show mean ± standard error of mean (SEM).

### ROI Analyses

We examined the effects of objective motion, pursuit, scene and their interactions using repeated measures ANOVAs in PPA, RSC and OPA.

In order to test for lateralization effects, we performed a 3x2x8 repeated measures ANOVA with the factors ROI, hemisphere and condition. There was no main effect of hemisphere (F (1, 12) = 0.027, p = 0.873), nor any interaction of hemisphere with any factor (hemisphere and ROI: F (2, 24) = 0.002, p = 0.998, hemisphere and condition: F (3.488, 41.855) = 0.25, p = 0.89 and hemisphere, ROI, condition: F (14, 168) = 1.618, p = 0.079). For the remaining analyses, we hence pooled data from both hemispheres for each ROI.

First, we conducted within-ROI analyses utilizing separate repeated measures ANOVAs with the factors scene, objective motion and pursuit for each ROI. We primarily focused on objective motion and retinal motion (i.e. the interaction between objective motion and pursuit), and on scene, and their interactions. These contrasts were completely balanced in terms of pursuit-related effects. We also report pursuit, but we note that it includes combined effects related to control of eye-movements, and to peripheral motion beyond the controlled visual screen induced by eye-movement, respectively.

### Motion in eye- and world centered reference frames

Figure 3 shows main effects related to motion in eye- and world-reference frames (i.e. retinal and objective motion) and to pursuit for each ROI. Each scene region had a distinctly different signature in terms of its motion response.

**Figure 3.**
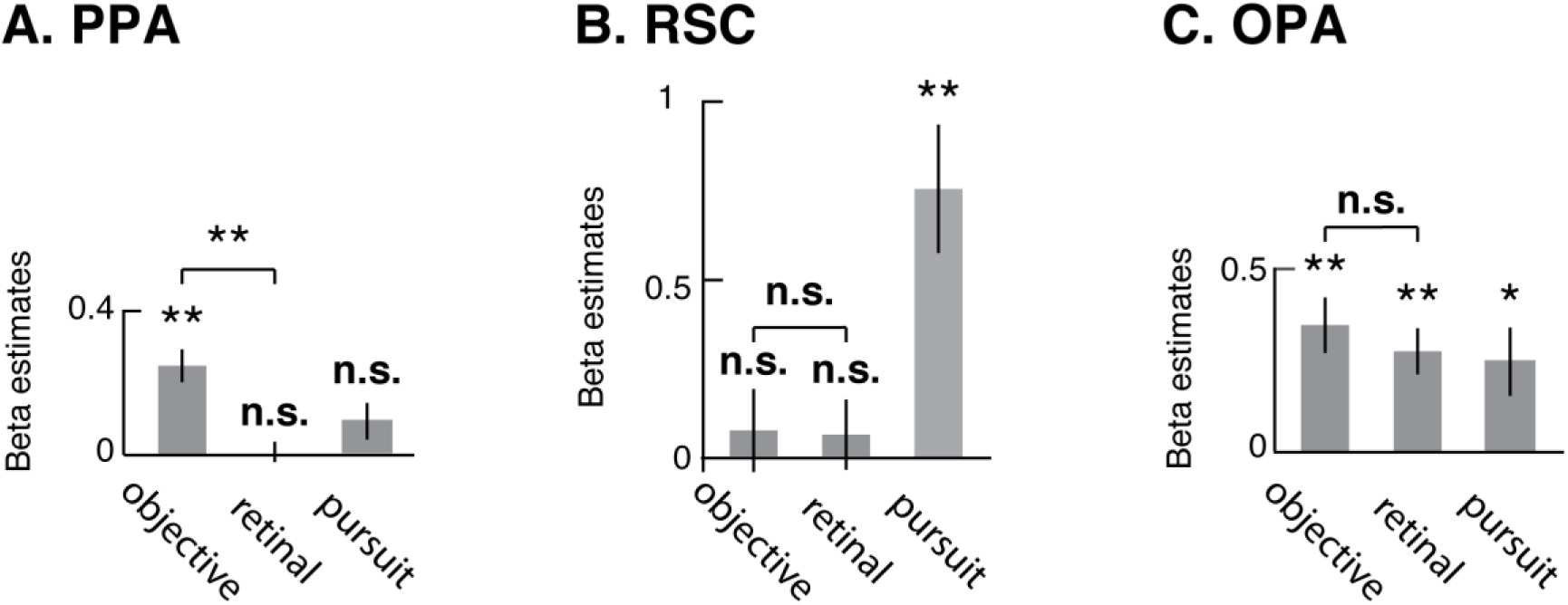
Responses of PPA, RSC, and OPA to objective motion, retinal motion, scene and pursuit. (A) PPA. (B) RSC (C) OPA. Note that data of each ROI was normalized. ** p < 0.001, *: p < 0.05. -values are Holm-Bonferroni corrected for 3 comparisons within each plot Plots show mean ± standard error of mean (SEM).

PPA responded robustly to objective motion (F (1, 33) = 34.56, p = 1*10^−6^), but not at all to retinal motion (F (1, 33) = 0.00047, p = 0.98), nor to pursuit (F (1, 33) = 3.19, p = 0.083). RSC did not respond to objective motion (F (1, 32) = 0.32, p = 0.57), nor to retinal motion (F (1, 32) = 0.41, p = 0.53), but it responded robustly to pursuit (F (1, 32) = 17.58, p = 0.0002). OPA responded to both, objective motion (F (1, 27) = 24.78, p = 0.00003), and retinal motion (F (1, 27) = 18.61, p = 0.00019), as well as to pursuit (F (1, 27) = 7.51, p = 0.011).

### Interactions between scene content and motion

Next, we tested whether motion responses were differentially driven by scenes and scramble content. This is important, as in principle some of the scene region’s motion responses could be accounted for by the increase in exposed scene-content by lateral motion, even though this was kept deliberately small in our stimuli, with 1.98 visual degrees in each direction out of 19 degrees stimulus width. The most important test was however whether objective motion responses persisted also for Fourier-scramble stimuli that lacked any scene content.

Figure 4A shows that all three ROIs responded robustly to scenes. Figure 4B shows that PPA responses were significant for objective motion, both during scenes (t (33) = 6.33, p = 1.1 * 10^−6^) and scrambled (t (33) = 3.51, p = 0.003), and that it had a significant interaction between objective motion and scene (t (33) = 2.57, p = =0.015). In RSC, neither scenes (t (32) = 0.96, p = 0.34) nor scrambled images (t (32) = 0.06, p = 0.95) resulted in significant responses during objective motion and there were no significant interactions (t (32) = 0.66, p = 0.51). In OPA, objective motion responses were significant both during scenes (t (27) = 4.15, p = 0.0009) and scrambled images (t (27) = 4.14, p = 0.0006), but there were no significant interactions (t (27) = 0.57, p =0.57). Figure 4C shows that PPA did not have retinal motion responses during scenes (t (33) = 1.3, p = 0.2) or scrambled images (t (33) = −1.47, p = 0.15) and there was no significant interaction (t (33) = 1.61, p = 0.12). In RSC, retinal motion responses were similar to PPA’s (during scenes: t (32) = 1.54, p = 0.13, during scrambled images: t (32) = −0.496, p = 0.62) and there was also no interaction between scene and retinal motion (t (32)=1.58, p = 0.12). In OPA, retinal motion responses were significant both during scenes (t (27) =3.39, p = 0.006) and scrambled images (t (27) = 3.46, p = 0.004), but there was no significant interaction (t (27) = 0.864, p = 0.395). P-values are Holm-Bonferroni corrected for 3 comparisons for each ROI. There were no significant pursuit-content interactions for any of the ROIs (p > 0.05).

**Figure 4.**
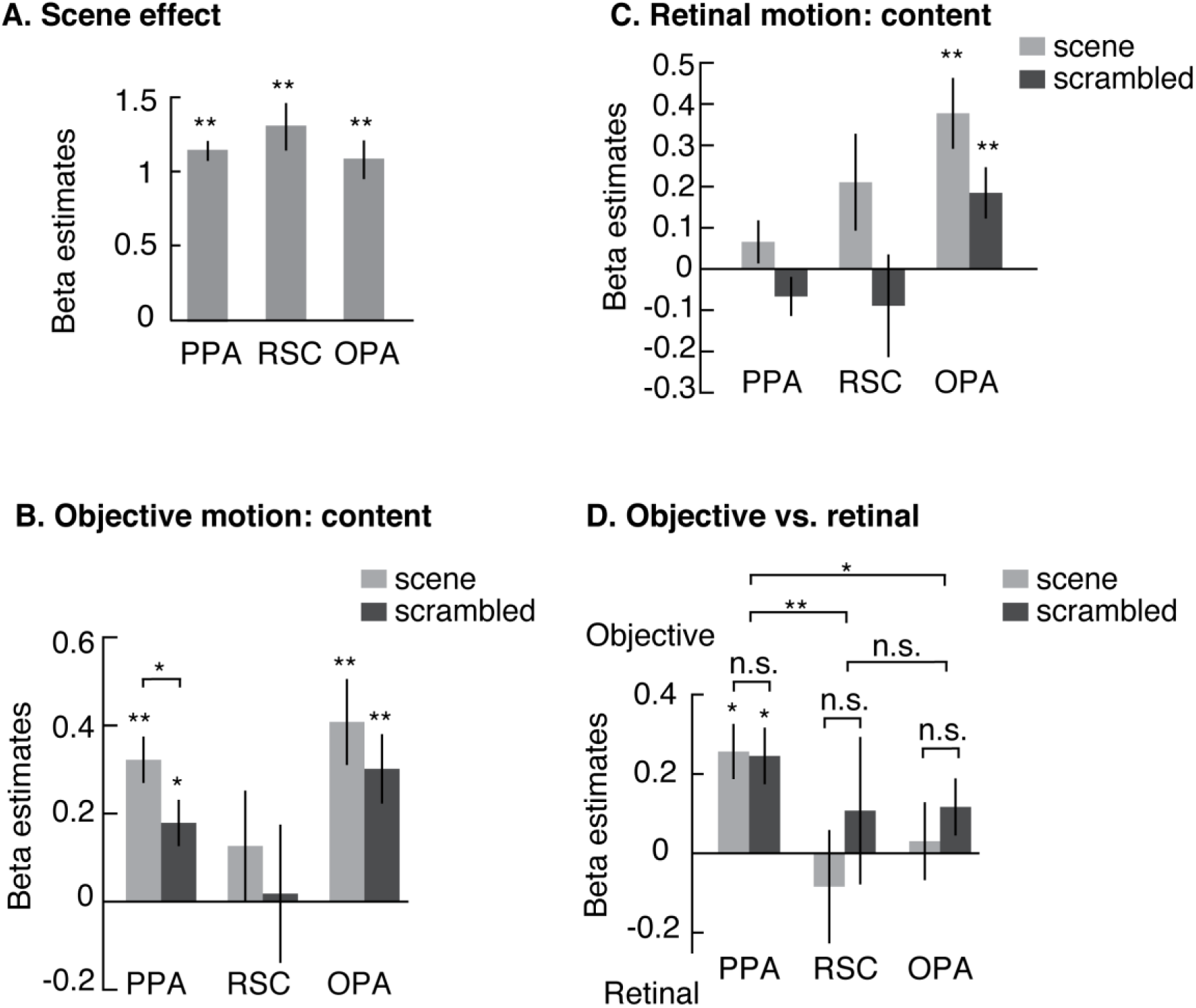
Effect of scene content on ROI responses. (A) Main effect of content, scene versus scrambled across all conditions (B) Responses to objective motion for scene and scrambled content. The figure shows responses to objective motion, separately for scene and scrambled backgrounds. (C) Responses to retinal motion for scene and scrambled content. The figure shows responses to retinal motion, separately for scene and scramble. (D) Objective versus retinal motion preferences across ROIs. Objective versus retinal motion responses, shown for scenes and scrambled backgrounds separately. Brackets across ROIs refer to tests calculated for the average response across scene and scramble *: p < 0.05, **: p < 0.001, Holm-Bonferroni corrected for 3 comparisons within each plot and each ROI.

Next, we tested which regions differed in the contrast (“objective” versus “retinal” motion). We analysed this contrast for scene and scrambled images separately. Figure 4D shows that PPA was the only region with a significant preference for objective motion both during scenes (t (33) = 3.79, p = 0.002) and scramble (t (33) = 3.52, p = 0.003). Neither RSC nor OPA had a preference for objective versus retinal motion (RSC: objective vs retinal during scenes: t (32) = −0.45, p = 0.65, objective vs retinal during scramble: t (32) = 0.38, p = 0.71, OPA: objective vs retinal during scenes t (27) = 0. 73, p = 0.47, objective vs retinal during scramble: t (27) = 1.36, p = 0.19). As shown in figure 4C, no region differed between scenes and scramble in their objective vs retinal motion responses (PPA: t (33) = 0.11, p = 0.91, RSC: t (32) = −0.67, p = 0.51, OPA: t (27) = −0.31, p = 0.76). P values are Holm-Bonferroni corrected for 3 comparisons for each ROI.

### Comparisons between ROIs

Next, we tested for differences between ROIs. We performed a 3 x 2 x 2 x 2 repeated measures ANOVA with the factors ROI (PPA, RSC, OPA), objective motion (on/off), pursuit (on/off) and scene (scene/ scrambled). The factor ROI had significant interactions with each of the remaining factors (objective motion: F (1.51, 40.71 = 13.53, p = 0.00013; retinal motion (i.e. objective motion and pursuit): F (1.62,43.79) =8.60, p = 0.001; scene: F (2,54) = 24.64, p = 2.5 * 10^−8^; pursuit: F (1.54,41.47) = 4.37, p = 0.027). There were no triple interactions. Direct comparisons between ROIs using Holm-Bonferroni corrected (for 3 comparisons) t-tests showed highly significant dissociations between ROIs. Figure 5 shows that PPA and OPA differed in their objective motion response from RSC (PPA vs. RSC (t (32) = 4.78, p = 0.0001); RSC vs. OPA (t (27) = −3.85, p = 0.002). PPA and OPA did not have any significant differences in their objective motion responses (t (27) = −0.86, p = 0.4). OPA differed in its retinal motion response from PPA (t (27) = 3.93, p = 0.003) and RSC (t (27) = 2.92, p = 0.01), whereas there was no difference between RSC and PPA in their retinal motion responses (t (32) = 0.68, p = 0.5). Only RSC and OPA responded to pursuit, with the former differing significantly from PPA (t (32) = −3.56, p = 0.0035).

**Figure 5.**
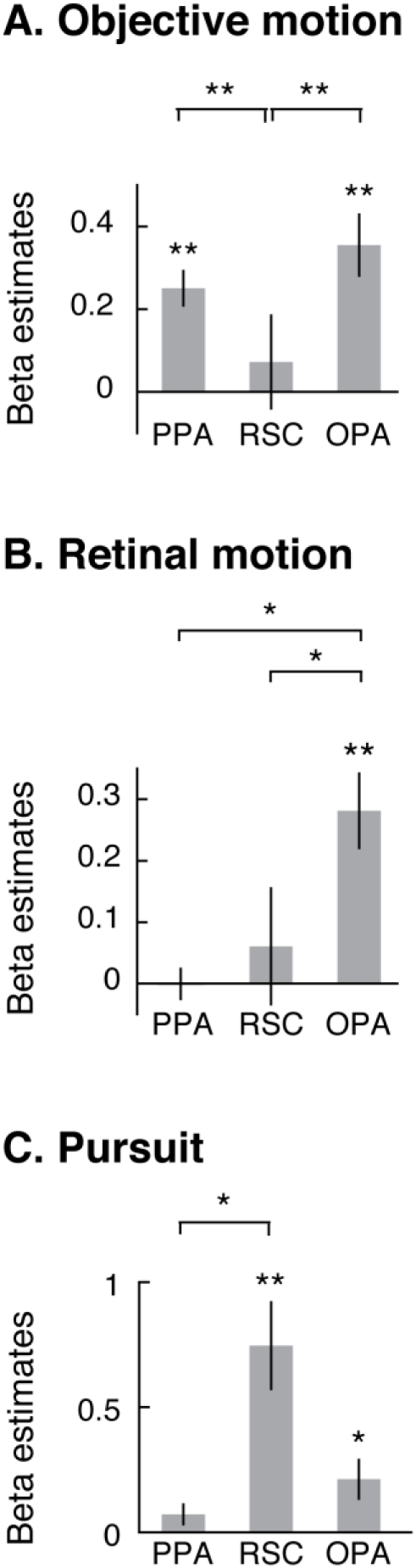
Responses of PPA, RSC, and OPA to objective motion, retinal motion and pursuit (A) Main effect of objective motion. (B) Main effect of retinal motion. Note that this is the interaction of objective motion and pursuit (C) Main effect of pursuit. Note that data of each ROI was normalized. **: p < 0.001, *: p < 0.05, Pairwise comparisons are Holm-Bonferroni corrected. Plots show mean ± standard error of mean (SEM).

Figure 4C shows that PPA had a higher preference to objective versus retinal motion than both, OPA (t (27) = 2.96, p = =0.01) and RSC (t (32) = 5.01, p = 0.57 * 10^−4^). There were no significant differences between OPA and RSC in their preference for objective vs retinal motion (t (27) = 1.3, p = 0.2).

In summary, objective motion responses were unique to PPA and OPA, and retinal motion responses were unique to OPA. PPA was the only ROI that showed significant objective motion preference over retinal motion. RSC stood out with strong pursuit responses, and PPA was the only region lacking them (Figure 5C).

With regards to content-motion interactions, PPA was the only region with content-motion interaction, with higher activation to motion during scenes than scramble. This interaction was not present in the comparison between objective motion with retinal motion. Importantly, the objective motion responses of PPA and OPA cannot be accounted for only by moving scene-content, as they were highly significant also for moving Fourier scramble.

### Adaptation Index

In a final analysis, we provide a comparison point to a related study. Golomb and colleagues used a similar paradigm yet with crucial differences. That study used saccades and still images rather than pursuit and continuous motion, respectively, and spatial updating (i.e. saccades) occurred in the absence of a stimulus (Golomb et al., 2011). All their results were quantified statistically in the form of adaptation indices, so we applied the same formula to our data to allow for a comparison. We did so for scene conditions only as they did not have scramble conditions, and we used the (purs/mov/sce), the condition with highest responses, as baseline. Using range-normalized data, we calculated the adaptation index for each ROI, each subject and each condition with scene background as follows:

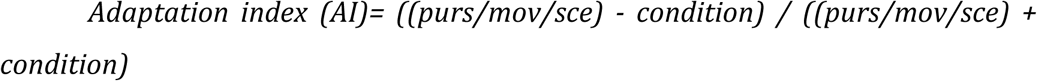

Figure 6 shows AIs for all ROIs and conditions. All p-values are Holm-Bonferroni corrected for 9 tests.

**Figure 6.**
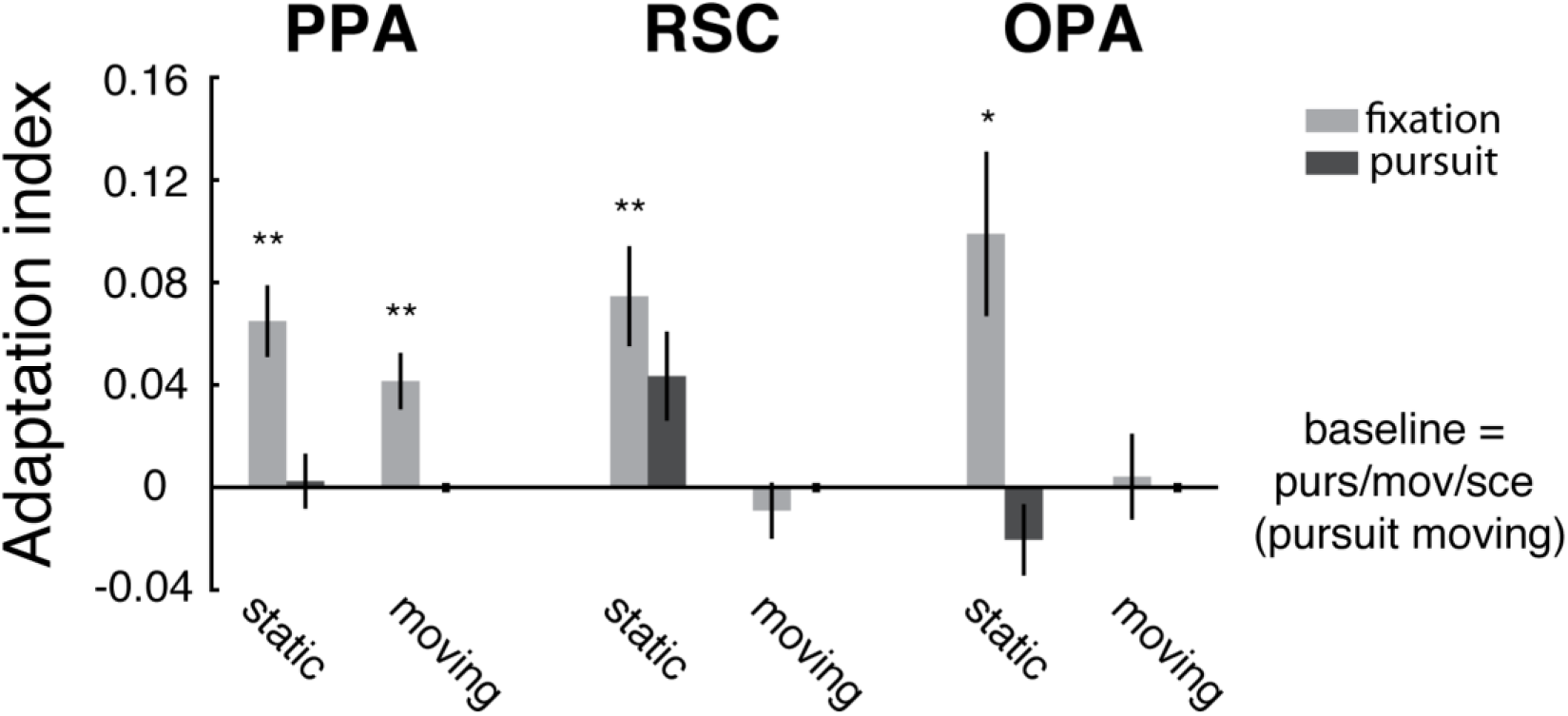
Adaptation indices for PPA, OPA and RSC. The baseline was calculated using (purs/mov) condition during scenes. Adaptation indices are only calculated during scenes. *: **=p < 0.05; **: p < <0.005, *=p,0.05 (Holm-Bonferroni corrected). Plots show mean ± standard error of mean (SEM).

Similar to the results of Golomb et al (Golomb et al., 2011), our results also revealed that PPA showed significant adaptation when eyes moved over static scenes but not when eyes were fixated and scene was moved. The (purs/sta/sce) condition (t (33) = 3.97, p = 0.003) showed significant adaptation whereas no adaptation was found for (fix/mov/sce) condition (t (33) = 0.24, p = 0.81). Moreover, our results reporting PPA responses during fixation and static background is also in agreement with their results in PPA during fixation. The (fix/sta/sce) condition (t (33) = 4.83, p = 0.0003) showed a significant adaptation. However, the main inconsistency between our and their results arises from PPA adaptation responses during eye movements. In our experiment, pursuit during scene motion during scenes (purs/mov/sce) resulted in highest beta responses and there was no adaptation whereas they reported significant adaptation during saccades on scrolling scenes.

Golomb et al (Golomb et al., 2011) reported that OPA showed adaptation during fixation on static scenes and during saccades on scrolling scenes and RSC showed no adaptation, noting that the responses they observed in OPA and RSC were much noisier than the responses observed in PPA. In OPA and RSC, our results indicate adaptation only during fixation and static background (fix/sta/sce) condition, and no adaptation was found for the other conditions (RSC: (fix/sta/sce):(t (31) = 3.91, p = 0.003), (fix/mov/sce): (t (31) = 2.56, p = 0.08), (purs/sta/sce): (t (31) = −0.87, p = 0.39); OPA: (fix/sta/sce): (t (21) = 3.08, p = 0.034), (fix/mov/sce): (t (21) = −1.52, p = 0.14), (purs/sta/sce): (t (21) = 0.26, p = 0.8)). These results reveal similarity in OPA, except that we again found opposite pattern during pursuit on moving scenes. In RSC, the only difference between our results and theirs is during fixation on static scenes.

### Whole brain Analyses

Beyond the ROI analyses, we performed additional random-effects analyses across all voxels of the brain in order to test whether other regions outside the selected ROIs responded to objective versus retinal motion contrast. Group level whole-brain analysis revealed that voxels overlapping with PPA showed activation at uncorrected levels (p < 0.05 uncorrected), as well as voxels near posterior occipital sulcus (POS), coinciding with motion sensitive region V3A (figure 6). Previously, V3A was shown to be sensitive to objective versus retinal motion contrast when random dots were used as stimuli (Fischer et al., 2012a). The activation overlapping with PPA has peak T-statistic values 3.26 and 1.90, respectively in right (30, −44, −6) and left PPA (−26, −48, −8).

**Figure 6.**
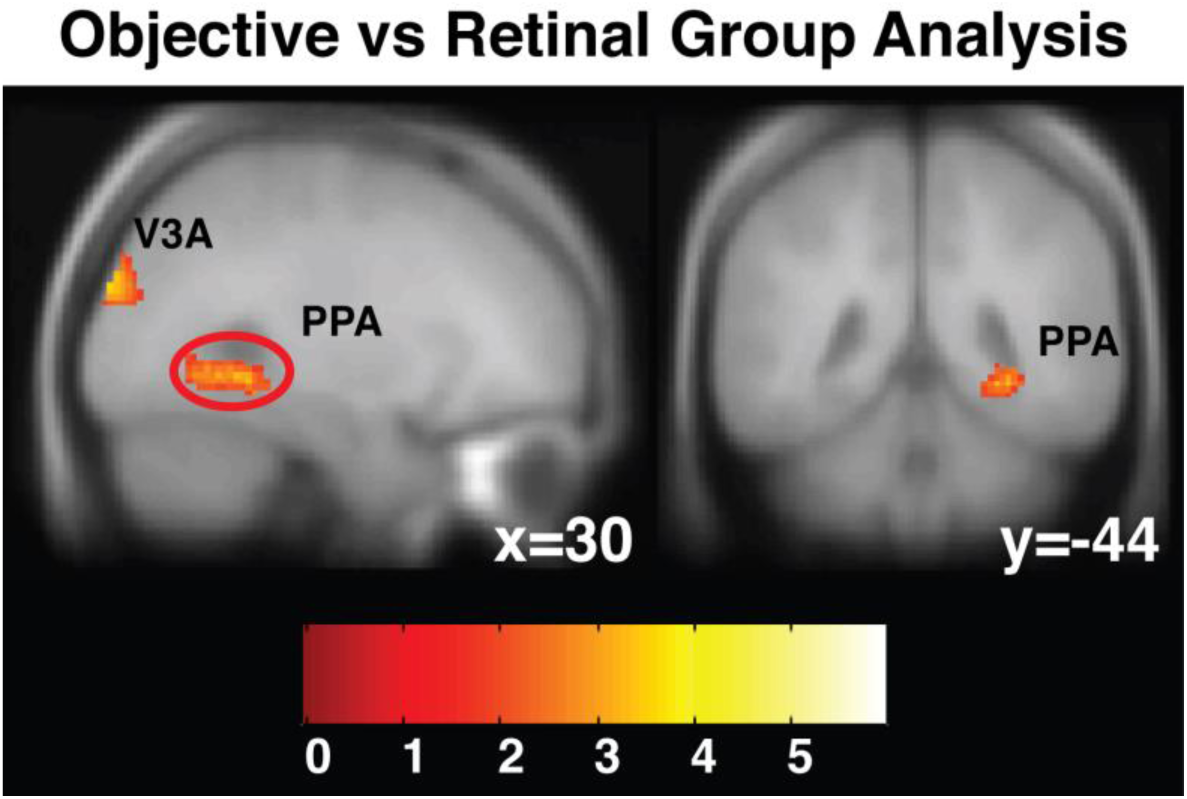
Objective versus retinal motion group data across all subjects. Voxels overlapping with PPA were activated at uncorrected thresholds, shown here inside the red circle in the sagittal slice in the left hemisphere and in the coronal slice in the right hemisphere. The dorsal activity near posterior occipital sulcus (POS) coincides with V3A (Fischer et al., 2012a). Peak t-statistic values 3.26 in right and 1.90 in left PPA (not shown here). For illustration only, the map is shown at p < <0.05 uncorrected.

### Behavioural Data

Participants performed a 1-back character-matching task intended to maintain vigilance and balance attention across conditions during the main experiment. The mean correct response rate was near ceiling with 0.85 ± 0.07 (mean ± S.D.). The mean response time was 0.57 ± 0.14 s (mean ± S.D.). We analysed response times using a 3-way repeated measures ANOVA with the factors objective motion, pursuit and scene. There were only significant effects of pursuit (F (1,16) = 5.06, p = 0.039) but no other main effects (objective motion: F (1,16) = 0.005, p = 0.95; scene: F (1,16) = 0.39, p = 0.54), nor interactions (objective and pursuit (i.e., retinal motion): F (1,16) = 0.93, p = 0.35; objective and scene: F (1,16) = 0.00029, p = 0.99; pursuit and scene (1,16) = 1.58, p = 0.23; objective, pursuit and scene: F (1,16) = 0.98, p = 0.34). There was also no difference between objective and retinal motion (t (1,16) = −0.81, p = 0.43).

### Eye tracking Data

We analysed eye-position error using the same 3-way ANOVA as used for the behavioural data. Like in behaviour, the only significant effect was the main effect of pursuit (F(1,67) = 682.38, p = 7.55 * 10^−37^), with no other main effects (objective motion: F(1,67) = 0.055, p = 0.82; scene: F(1,67) = 0.00014, p = 0.99), or interactions (retinal motion: F(1,67) = 0.28, p = 0.60; objective motion and scene: F(1,67) = 0.14, p = 0.71; pursuit and scene: F(1,67) = 2.80, p = 0.10; objective motion, pursuit and scene: F (1,67) = 0.16, p = 0.69). There was no objective versus retinal difference (t (67) = 0.13, p = 0.90).

## Discussion

The aim of this study was to determine to which extent scene responsive regions PPA (Aguirre et al., 1998; Epstein and Kanwisher, 1998), RSC (Maguire, 2001) and OPA (Dilks et al., 2013; Grill-Spector, 2003; Hasson et al., 2003; Nakamura et al., 2000) encode visual motion cues in eye-centered (i.e. retinal) or world-centered (i.e. objective) reference frames. We used a previously established paradigm that combined pursuit eye-movements with on-screen motion (Fischer et al., 2012a) to determine retinal- and objective motion responses in independently defined scene-selective ROIs. To dissociate scene-dependent effects from low-level motion responses, we repeated all conditions using natural scenes and their Fourier scrambled versions, allowing examining motion-content interactions.

We found that each of the scene responsive regions had unique and dissociable response profiles. In particular, PPA was unique in its sensitivity to objective motion while not responding at all to retinal motion or to pursuit eye movements. While PPA’s response to objective motion was enhanced by scene-content, its invariance to retinal motion and its response to objective motion were both preserved also for scrambled content. In contrast, OPA was the only scene region responsive to retinal motion (along with objective motion), and RSC responded strongly to pursuit but not to either motion type. Apart from providing a functional segregation of scene regions in terms of motion reference frames, the present results show that PPA is able to represent a stable environment during self-induced retinal motion.

### Controversy over ventral stream reference frame

The visual system uses different spatial representations in order to keep a stable vision of the world and these representations are reported in various brain areas. For instance, retinotopic representations are reported in early visual areas as well as higher level visual regions (Gardner et al., 2008; Golomb and Kanwisher, 2012) whereas world-centered spatial representations are thought to be encoded in dorsal regions, parietal regions and high level motion areas (Crespi et al., 2011; d’Avossa et al., 2007; Fischer et al., 2012a, b; Galletti et al., 1993; Ilg et al., 2004). The spatial representation in ventral regions is rather under debate; both retinotopic (Amano et al., 2009; Larsson and Heeger, 2006) and spatiotopic (McKyton and Zohary, 2007) maps are reported in LOC (lateral occipital cortex) and previous research indicate both eye centered (Golomb et al., 2011; Ward et al., 2010) and world centered (MacEvoy and Epstein, 2007) spatial representation in scene selective parahippocampal cortex.

### Choice of reference frame in scene regions

Previous studies demonstrated viewpoint dependency and specificity in scene representation of PPA in context of static scene views (Epstein et al., 2003; Epstein et al., 2007; MacEvoy and Epstein, 2007; Park and Chun, 2009; Park et al., 2010). The objective motion contrast used here is – conceptually – comparable to the kind of viewpoint changes of static scene views used in previous studies, since both involved a gradual, even if step-wise, lateral translation of the scene. It is worth noting that none of the previous studies mentioned involved motion or pursuit eye movements, since they were mostly carried out during eye fixation. The problem lies within the never changing position of eyes. When the eyes are fixated, the motion of the scenes in the background results in a combined objective and retinal motion. Therefore, it is not possible to differentiate whether the mechanism behind the mentioned PPA responses, which was described as viewpoint dependency or specificity, are retinal or world-centered viewpoint dependency. In their 2011 study, Golomb and her colleagues tried to address this problem and they found evidence for both reference frames, but concluded that the results in PPA are mainly driven by retinotopic responses (Golomb et al., 2011). Using an adaptation paradigm, they reported that PPA was adapting to successive views of the same scene when the participants did saccades across the scenes, which resulted in matching retinal input (Golomb et al., 2011). We calculated adaptation indices similar to theirs to directly compare our results. Since our experimental design did not include a novel scene condition, we used purs/mov/sce condition as baseline, which combines smooth pursuit eye movements on a moving background during scenes. This was feasible because this condition covered both types of motion (smooth pursuit eye movements and objective motion) and elicited the highest beta responses in PPA. Although our results were similar with theirs during most conditions, one striking difference is the opposite pattern of results during pursuit eye movements on moving background during scenes ((purs/mov/sce) condition). Unlike their results, we did not observe any adaptation during this condition.

The difference between our results and the results of Golomb et al (Golomb et al., 2011) might be rising from the use of saccades in their study versus the use of smooth pursuit eye movements in our experiment. Although smooth pursuit and saccades are traditionally thought to have partially overlapping neural networks, it is known that these networks are not identical. For instance, smooth pursuit eye movements and saccades are processed in different sub-regions of frontal eye fields (Rosano et al., 2002). Moreover, pursuit related neurons are found in regions such as V5/MT, MST and VIP (Ilg, 2008; Thier and Ilg, 2005). V3A is also thought to have a role in encoding pursuit (Fischer et al., 2012a), while it does not have any role in encoding saccades. Additionally, saccadic suppression generates stable visual input by actively suppressing the vision, particularly via suppression of magnocellular pathway (Burr et al., 1994; Thiele et al., 2002). Thus, it is possible that saccadic eye movements and smooth pursuit eye movements engage different brain regions.

Another difference is that in their experiment (Golomb et al., 2011), the saccades were executed during blank screen, preventing remapping of the actual scene, whereas in our experiment smooth pursuit eye movements were carried out on scenes, allowing for direct continuous remapping. Lastly, it is also possible that using saccades and pursuit eye movements during the stimuli might be resulting in perceptually different inputs. Further studies are needed in order to directly compare the effect of eye movement types during dynamic scene perception.

It is interesting to note that in the present study, similar to the results of Golomb et al (Golomb et al., 2011), RSC did not adapt to overlapping scenes when eyes fixated whereas this condition was previously reported to result in significant adaptation in RSC (Park and Chun, 2009). As they discussed, this difference could be driven by different tasks used in our experiment and previous studies(Golomb et al., 2011).

### Similarity to V3A

The objective motion preference of PPA was so pronounced, also with regard to the complete absence of retinal motion, that it bears similarity to that of dorsal motion areas V3A and V6 that have a similar response profile (Fischer et al., 2012a).

### Content – motion interactions

The interaction of PPA responses between objective motion and scene content, in that it showed higher responses to motion during scenes than during scrambled images, extend our previous findings, which were carried out by moving scenes and scrambled images during fixation (Korkmaz Hacialihafiz and Bartels, 2015). One could argue that the objective motion response in PPA could be explained by the fact that objective motion conditions revealed more of the scene image than static conditions. This is however an unlikely explanation. First, the preference for objective motion was still significant in PPA in context of scrambled images lacking any scene content. Second, the *difference* contrast of (objective – retinal) was equally large for scene and for scramble, implying that content played no role whatsoever here. Third, the displacement of the background images was a mere 1.98 visual degrees in either direction, with a field of view of 19 degrees, hence only revealing marginally new scene content, in the periphery where it is hard to recognize. It is thus highly unlikely that motion responses in PPA can be explained by increased scene content only.

### Responses in OPA

OPA was the only scene responsive region that showed significant retinal motion responses, and could be differentiated by this response profile from PPA and RSC, which lacked retinal motion responses entirely. OPA responded to both objective and retinal motion equally. These findings, and the fact that OPA lacks adaptation during both retinal and objective motion, are compatible with the previously proposed theory that OPA constitutes a comparably early level in the hierarchy amongst scene responsive regions due to its physical location and close proximity to dorsal regions (Dilks et al., 2011; Dilks et al., 2013; MacEvoy and Epstein, 2007). This view was further supported by a number of studies reporting low-level features of OPA. For instance, OPA has been shown to have lower visual field bias (Silson et al., 2015) and its receptive field sizes were smaller compared to other scene responsive ROIs (MacEvoy and Epstein, 2007; Silson et al., 2015).

### No pursuit in PPA

Another notable result is that PPA did not respond to pursuit at all, whereas OPA and RSC did. This shows an extreme degree of invariance to motion in the peripheral field of view invariantly induced by pursuit, and potentially powerful mechanisms cancelling eye-movement related jitter everywhere in its visual representations. Overall, this is compatible with the view of PPA representing scenes (and any visual signal) in a manner robustly independent of precise fixation points. This, together with PPA’s unique representation of motion only in world- but not eye-centered coordinates would clearly facilitate perception of a continuous world from discontinuous views. Stable scene representations are also supported by the ability to extrapolate views through boundary extension, a property PPA shares with RSC (Park et al., 2007).

### Responses in RSC

The lack of motion responses in RSC is comparable to previous studies reporting that RSC has viewpoint invariant properties (Park and Chun, 2009; Park et al., 2010) and that RSC shows little change in activity when the stimuli presented in different locations (other than central fixation point) (MacEvoy and Epstein, 2007; Ward et al., 2010). However, the pursuit response in RSC, especially when considered together with the lack of motion responses, is rather interesting. Previously, correlations between the activity of frontal eye fields (FEF) and Brodmann areas that constitute retrosplenial cortex were found during resting state (Hutchison et al., 2012). In primates, retrosplenial cortex is found to have projections to peripheral vision regions in MT and MST (Palmer and Rosa, 2006). Moreover, a recent fMRI study in humans showed that in comparison to other scene regions PPA and OPA, RSC showed more peripheral bias (Baldassano et al., 2016) and RSC is located near the peripheral visual regions in V1 and V2 (Nasr et al., 2011). During pursuit condition, peripheral visual changes are not controlled in the present study. Thus, pursuit responses in RSC could have been driven by peripheral bias in scene representation in RSC. More studies are needed in order to understand the mechanisms behind the pursuit related responses in RSC.

## Conclusion

In conclusion, our results reveal a novel dissociation between scene responsive regions PPA, OPA and RSC in their responses to real-world motion. In particular, we found that PPA could be differentiated from OPA and RSC in objective motion preference to retinal motion of PPA and no preference in OPA and RSC. OPA, given its location and probably mid-level position in scene processing hierarchy, was responsive to both objective and retinal motion. RSC, which is thought to be involved in higher-level functions such as spatial navigation and which was shown to have viewpoint independent responses, lacked motion responses but was activated by smooth pursuit eye movements. Altogether, these findings shed light into our understanding of how PPA, OPA and RSC responses are adjusted by real world motion. Present findings suggest a role for PPA in providing input for stable visual perception, by separating real-world motion from self-induced retinal motion, similar to the selectivity previously shown in V3A and partially in V6 (Fischer et al., 2012a).

## Acknowledgements

This work was funded by the Centre for Integrative Neuroscience Tübingen through the German Excellence Initiative (EXC307) and by the Max Planck Society, Germany.

